# Towards increased accuracy and reproducibility in SARS-CoV-2 next generation sequence analysis for public health surveillance

**DOI:** 10.1101/2022.11.03.515010

**Authors:** Ryan Connor, David A. Yarmosh, Wolfgang Maier, Migun Shakya, Ross Martin, Rebecca Bradford, J. Rodney Brister, Patrick SG Chain, Courtney A. Copeland, Julia di Iulio, Bin Hu, Philip Ebert, Jonathan Gunti, Yumi Jin, Kenneth S. Katz, Andrey Kochergin, Tré LaRosa, Jiani Li, Po-E Li, Chien-Chi Lo, Sujatha Rashid, Evguenia S. Maiorova, Chunlin Xiao, Vadim Zalunin, Kim D. Pruitt

## Abstract

During the COVID-19 pandemic, SARS-CoV-2 surveillance efforts integrated genome sequencing of clinical samples to identify emergent viral variants and to support rapid experimental examination of genome-informed vaccine and therapeutic designs. Given the broad range of methods applied to generate new viral genomes, it is critical that consensus and variant calling tools yield consistent results across disparate pipelines. Here we examine the impact of sequencing technologies (Illumina and Oxford Nanopore) and 7 different downstream bioinformatic protocols on SARS-CoV-2 variant calling as part of the NIH Accelerating COVID-19 Therapeutic Interventions and Vaccines (ACTIV) Tracking Resistance and Coronavirus Evolution (TRACE) initiative, a public-private partnership established to address the COVID-19 outbreak. Our results indicate that bioinformatic workflows can yield consensus genomes with different single nucleotide polymorphisms, insertions, and/or deletions even when using the same raw sequence input datasets. We introduce the use of a specific suite of parameters and protocols that greatly improves the agreement among pipelines developed by diverse organizations. Such consistency among bioinformatic pipelines is fundamental to SARS-CoV-2 and future pathogen surveillance efforts. The application of analysis standards is necessary to more accurately document phylogenomic trends and support data-driven public health responses.

## Introduction

The COVID-19 pandemic stimulated an unrivaled level of research in a short period of time resulting in an unprecedented amount of public genomic data for any single taxon – SARS-CoV-2. Genomes inferred from Next-Generation Sequencing (NGS) data enable global tracking of transmission routes and detection of new SARS-CoV-2 strains that can affect the efficacy of therapeutics or molecular diagnostic assays. Analysis of prevalent genomes supports the design of plasmids and pseudoviruses to perform experimental testing, such as neutralization assays to determine which monoclonal antibodies retain efficacy. The CDC provides viral strains that are used by the FDA, among other organizations and purposes, to guide which therapeutics receive emergency use authorization per each region (https://www.cdc.gov/coronavirus/2019-ncov/lab/grows-virus-cell-culture.html). In addition, the FDA requires submission of NGS data and the interpretation by pharmaceutical companies for new drug applications. Therefore, accurate and reproducible detection of mutations in SARS-CoV-2 sequencing data is critical for effective management of the SARS-CoV-2 pandemic (https://covid.cdc.gov/covid-data-tracker/#variant-proportions), and the development of vaccines and antiviral drugs.

As of August 17 2022, there have been over 11.7 million SARS-CoV-2 sequence submissions from over 7.9 million samples submitted to the National Institute of Health (NIH), National Library of Medicine (NLM), National Center for Biotechnology Information (NCBI) open access repositories GenBank^1^ and the Sequence Read Archive (SRA)^2^; of these, data was submitted to both GenBank and SRA for over 3.8 million samples. This sequence data represents the culmination of efforts across many different research institutions and public health laboratories around the world, using different library preparation methods, sequencing technologies and analysis methods. SARS-CoV-2 samples have been prepared in multiple different ways for sequencing, including shotgun, capture based methods, and the commonly-used tiled amplicon-based approaches, which continue to evolve. These have been coupled with the full complement of today’s most popular sequencing technologies including short read Illumina sequencers and long read sequencers like Oxford Nanopore Technologies (ONT) and Pacific Biosciences (PacBio). Bioinformatic analysis pipelines to generate SARS-CoV-2 genomes and to identify sequence variations also differ widely, even within a specific methodology/technology, with many developed and optimized for entirely different use cases. Therefore, no worldwide standards exist for generating SARS-CoV-2 genomes from NGS data, or for identifying important sequence variation therein.

Consensus-based approaches using NGS data have been a standard practice for SARS-CoV-2 mutation discovery, and a significant number of viral consensus sequences have been made available in open access or restricted access repositories such as GenBank and GISAID (https://www.gisaid.org/) to support public health research. Reference alignment-based variant discovery using NGS data is not new, and has been widely used in human population studies. The performance of such approaches using different processing pipelines, including read aligners, variant callers, and sequencing instruments, has been assessed by various studies^3–9^ resulting in best practices for germline^10^ and somatic variant detections.^11^ However, the performance of NGS pipelines for SARS-CoV-2 mutation detection has not been systematically assessed. Such evaluation is urgently needed with the on-going SARS-CoV-2 pandemic, as it will provide guidelines and best practices specific to SARS-Cov-2 and mutation detection by research groups, clinical sequencing labs and public health agencies,^12^ and set the stage for similar types of analyses in the future.

To address this issue, as part of the Foundation for the National Institutes of Health’s (FNIH) Accelerating COVID-19 Therapeutic Interventions and Vaccines Tracking Resistance and Coronavirus Evolution (ACTIV TRACE) initiative (https://www.nih.gov/research-training/medical-research-initiatives/activ/tracking-resistance-coronavirus-evolution-trace), seven groups across government, industry, and academia assessed a number of different approaches to SARS-CoV-2 NGS data analysis. The pipelines considered here were initially developed for each participating groups’ specific needs and are used here to analyze a representative subset of data available from the SRA. Thus, while some differences in results were expected, given that they all aim at calling variants for SARS-CoV-2 data, it is reasonable to expect that, given the same input, they should yield overall comparable results. Comparing results across the pipelines using common input datasets helped identify unexpected differences in analysis results and determine standardized protocols to improve the consistency of results across pipelines. We describe below our analysis of raw sequencing data and the development of quality assurance protocols to identify the most supported sequence variation calls from a sequenced sample, which in turn can be used for downstream ACTIV TRACE efforts to assess therapeutic efficacy in response to the evolving SARS-CoV-2 virus.

## Results

### SARS-CoV-2 consensus bioinformatic workflow overview

With the great global investment in sequencing the SARS-CoV-2 virus using a broad array of disparate techniques, we aimed to characterize bioinformatic methods for identifying variants from NGS sequencing data to support accurate and reliable use of these data in downstream analyses such as for phylogenomic and pharmaceutical applications. When the work was initiated, Illumina platform data was the most prevalent data type, with Oxford Nanopore Technologies (ONT hereafter) data the next most common, hence the current focus. Currently, however, PacBio data is approximately as common as ONT. The generic bioinformatics workflow for processing SARS-CoV-2 genomes has several steps as outlined in **Figure 1**. These include data retrieval, de-hosting, read clean-up (QC), alignment, variant calling, and variant filtering (post-processing). We evaluated 7 bioinformatics workflows which include many of the most commonly used tools and software packages for each step of the bioinformatics process (QC, alignment, variant calling). A more detailed comparison of each pipeline’s software and parameters is summarized in **Supplemental Table 1**. Of note, in agreement with the variety of software available for each, we found a much greater diversity in software used for Illumina platform data (**Figure 1A**) than ONT (**Figure 1B**), further supporting the generalizability of our results. To perform this analysis, a preliminary dataset, termed “Dataset 1,” was used to develop each pipeline’s protocol. A subsequent dataset, the “Dataset 2,” was used to compare concordance across pipelines.

**Figure 1.**
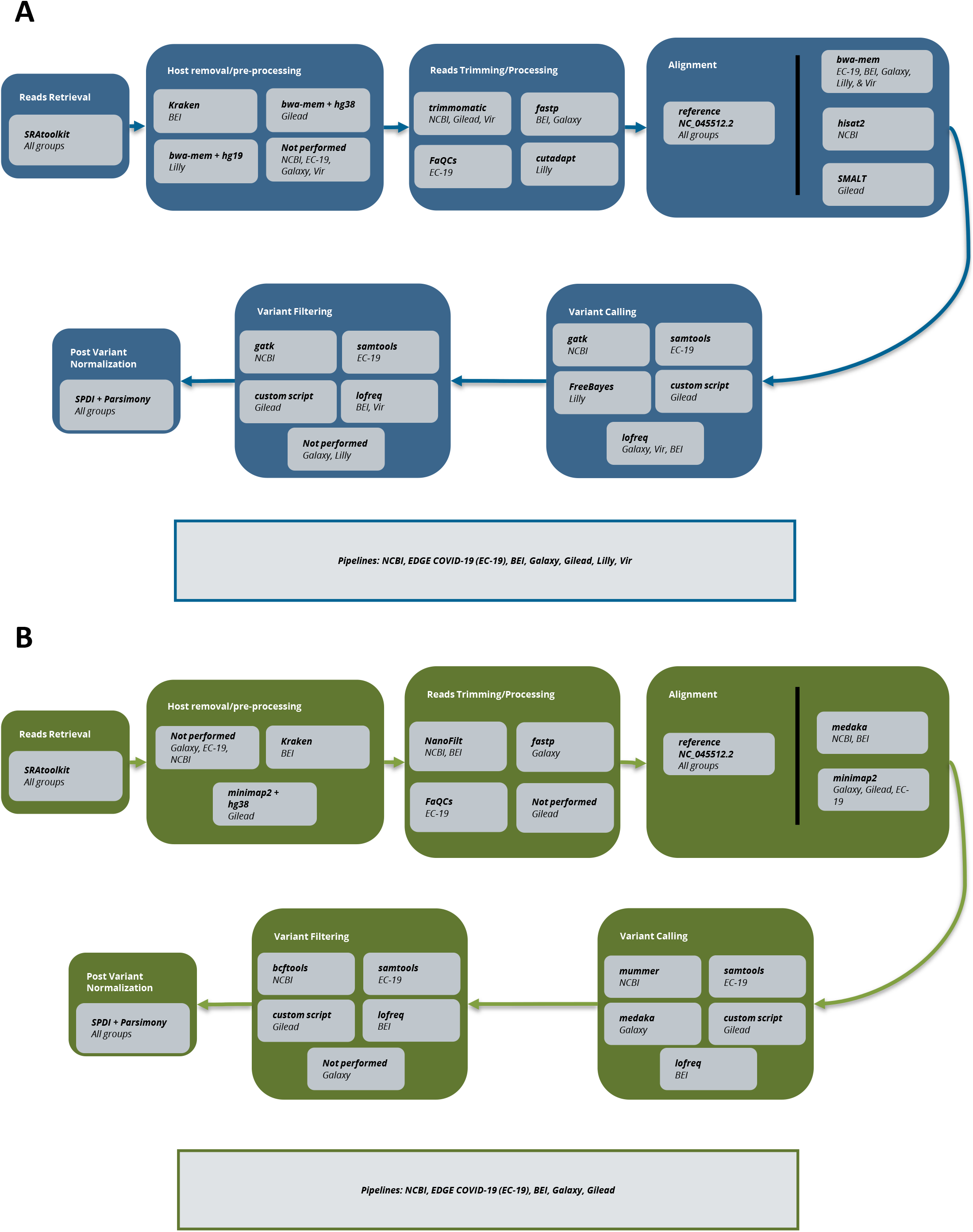
Analysis schematic. A) Illumina Platform Variant Calling. B) Oxford Nanopore Technologies (ONT) Variant Calling. For each sequencing technology, the main steps of variant calling are broken out in large boxes: read retrieval, host removal, read trimming, alignment, variant calling, variant filtering, variant normalization. For each step, software used by each group/s pipeline is noted.

### Data pre-processing impacts on variant calling

Initial attempts at comparison across pipelines identified large differences between the pipelines, more than was expected. To ensure these differences were genuine, we first examined differences in how the pipelines pre-processed the data. First, it was noted that not all groups were taking steps to exclude possible host-read contamination. As shown in **Figure 2A**, failure to remove host-read data can result in spurious calls, and thus artificially reduce the observed frequency of true calls. Additionally, differences in how workflows handled primer trimming were found. As seen in **Figure 2B**, failure to trim primer sequence from reads results in a reduction in the reported frequency of many variants in primer binding sites. This can reduce variant frequency below a threshold for consideration, 15% for the purposes of the work here, or below the threshold for incorporation into a consensus sequence (50%, e.g.). Unfortunately, accounting for primer sequence, especially for reanalysis of SRA data, can be tedious as such information is typically not available.

**Figure 2.**
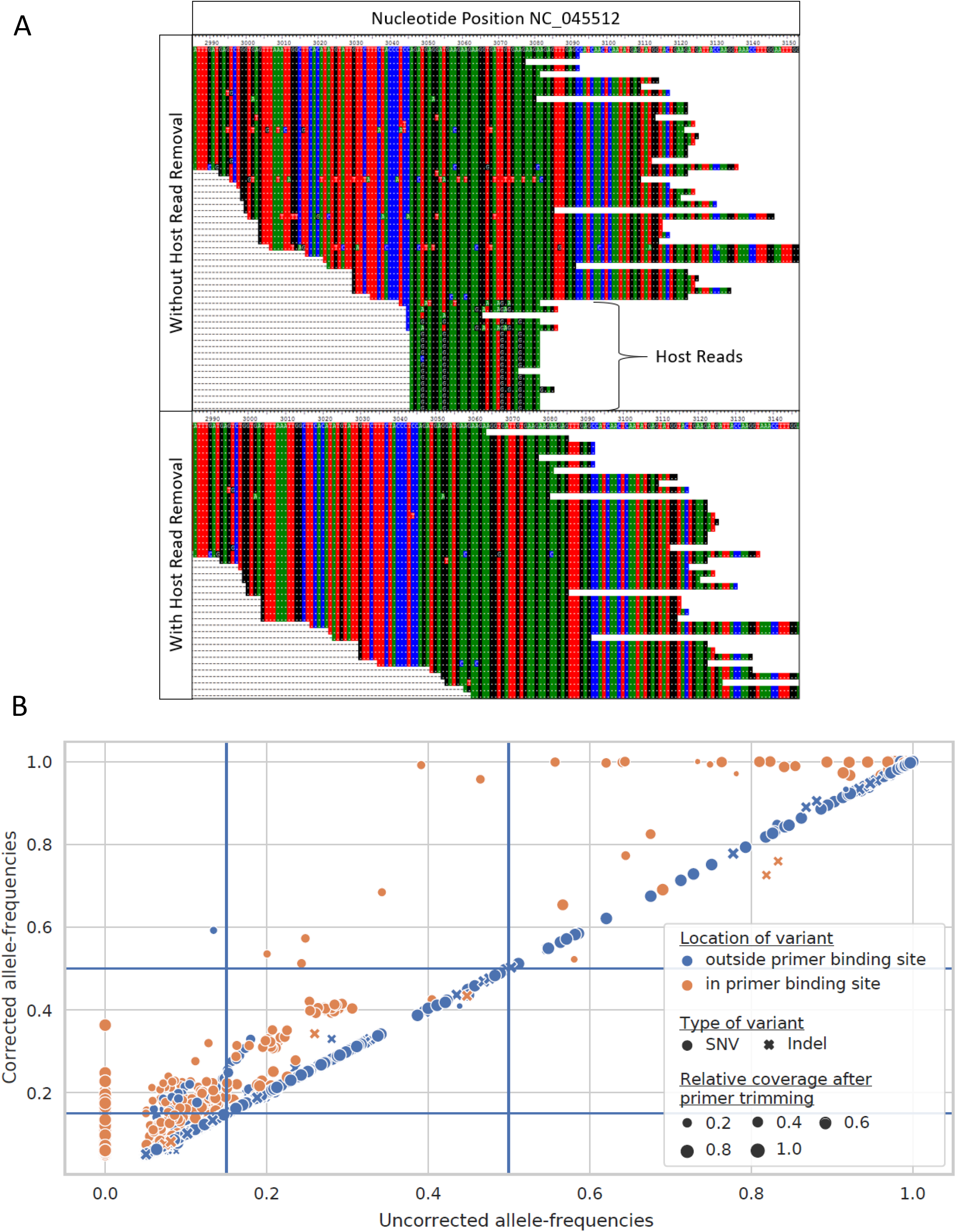
Impact of read cleanup. A) Impact of de-hosting on read alignment. Removal of host reads from RNAseq SARS-CoV-2 sequencing result, SRA run SRR12245095, reduced the potential for false positive variant calls. In the top panel, additional mutations were present in aligned reads between positions 3049-3076 of NC_045512 when host reads were not removed. After excluding host reads (bottom panel), reads containing the mutations were no longer observed B) Impact of primer trimming on variant calls. Allele-frequencies of variants called after trimming primer sequences from aligned reads (corrected allele-frequencies) are plotted against allele-frequencies of the same variants called without primer trimming (uncorrected allele-frequencies). Primer trimming increases called allele-frequencies of most within-primer binding sites variants. Blue lines represent the allele-frequency thresholds used in this study to filter variant calls (AF >= 0.15) and to call consensus variants (AF >= 0.5). Primer trimming lifts the allele-frequencies of 61 within-primer binding site variants above the threshold for retaining them, and enables calling of six additional consensus variants.

### A parsimony normalization method to standardize variant reporting across all workflows allowed vis-a-vis comparison of variants

Comparison of results was further complicated by differences in how variants are reported, with this issue most pronounced in the case of Insertions and Deletions (InDels). To address this, two steps were taken. First the SPDI algorithm^13^, used as part of the dbSNP and ClinVar databases at NCBI, for human genetic variants was adapted for use with SARS-CoV-2. While this greatly reduced the number of observed discrepancies in formatting, a number of issues with InDels remained. To this end, a Parsimony algorithm was developed (see Methods) to ensure the number of preceding and subsequent bases around an event were consistent, and that the events were left-aligned to a common starting position. Together these approaches help minimize the possibility that the remaining differences are artifactual, and thus increase the likelihood that they represent genuine differences between pipeline results.

### Allele Frequency and Depth of Coverage are important parameters and determine consistency of variant calls across workflows

To assess parameters that impact agreement between pipelines and across platforms a Receiver-Operating Characteristic (ROC) analysis was conducted. For this purpose, variants identified by greater than 5 workflows (or by both technologies) were considered a True Positive while calls made by fewer workflows (or a single technology) were considered a False Positive. The rationale is that if a result is accurate, it should be found by any pipeline (or technology), regardless of the implementation specifics, though this approach cannot guarantee the agreed upon results are accurate. Additionally, this means that the ROC AUCs cannot be directly compared between groups. Looking at the ROC curves to identify the inflection point across parameter settings can suggest settings that maximize True Positive results, across workflows or technologies, while minimizing False Positive results. **Figure 3A+B** and **Figure 3C+D** show the results for comparing results across pipelines for Illumina and ONT platform results respectively, while **Figure 3E+F** and **Figure 3G+H** show the results for comparing between technologies for Illumina and ONT platform results respectively. The most sensitive discriminator, the parameter for which a sharp inflection point was identified for most workflows, for results across pipelines was found to be Alternate Allele Depth (ALTDP), **Figure 3A+C+E+G**, while the best discriminator for results between technologies was found to be Alternate Allele Frequency (AF), as depicted in **Figure 3B+D+F+H**. Accordingly, a minimum ALTDP of 50 and a minimum AF of 50% was used for subsequent analyses. Additionally, as overall read depth (DP) was also found to perform well for comparison of results across pipelines (data not shown) a minimum DP of 100 was used as well. Additionally, records for which less than 50% of the reference sequence was covered or for which the average coverage was less than 100 were excluded, and as not all workflows supported the analysis of single-end read data, these were excluded as well. Finally, as many of the InDel discrepancies that could not be resolved with the combination of SPDI and Parsimony approaches were found to be in homopolymer regions, these were excluded as well (**Supplemental Figure 1**).

**Figure 3.**
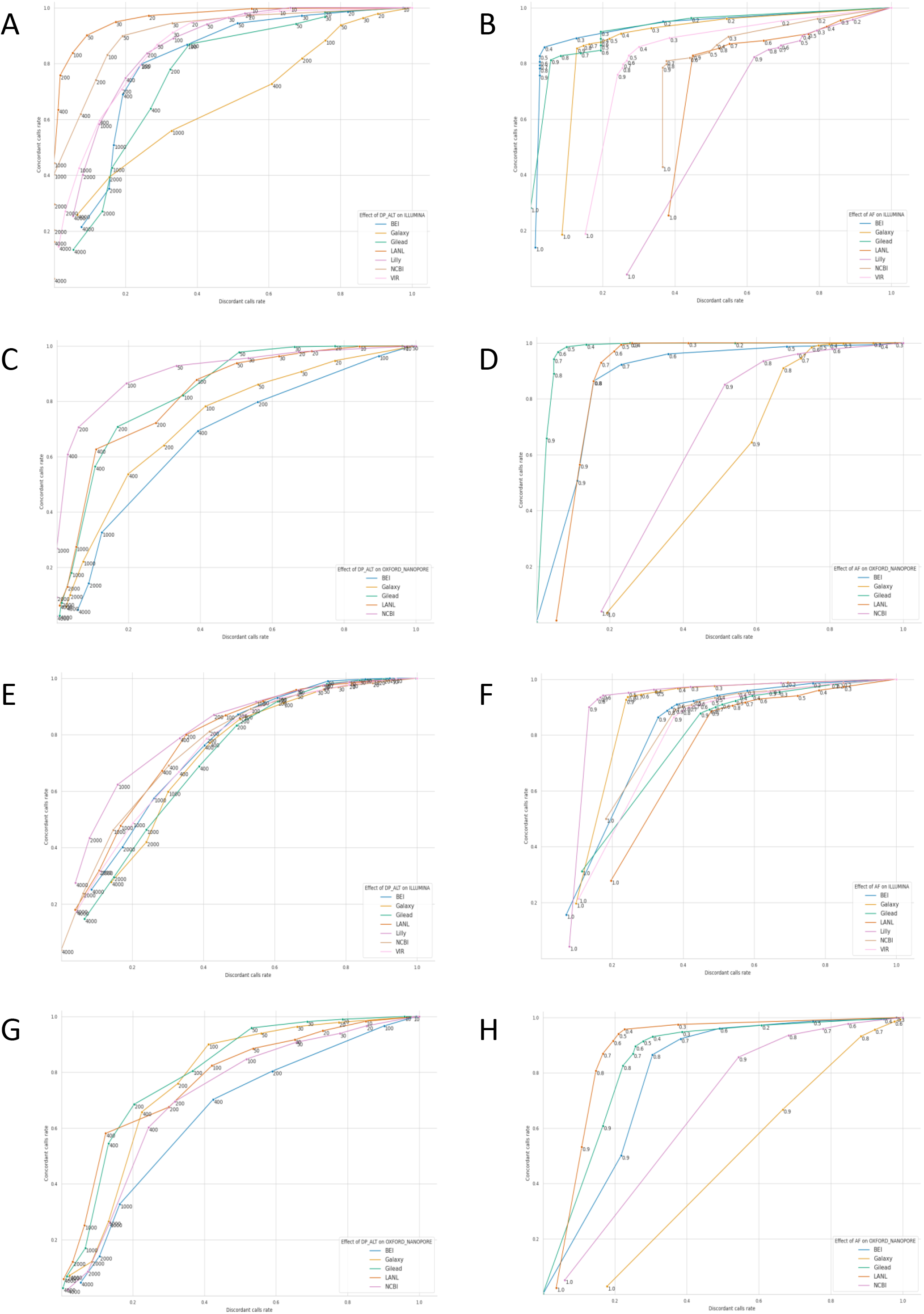
Identification of filtering parameters inflection points. A+C) Effect of alternate allele read depth (AltDP) on Accuracy and Specificity of each pipeline with regards to calls made by the majority of groups. B+D) Effect of alternate allele frequency (AF) on Accuracy and Specificity of each pipeline with regards to calls made by the majority of groups. A+B) Impact of AltDP and AF on pipeline accuracy and specificity for Illumina pipelines. C+D) Impact of AltDP and AF on pipeline accuracy and specificity for Illumina pipelines. E+F) Impact of AltDP on accuracy and specificity of calls made by both platforms for Illumina pipelines. G+H) Impact of AltDP on accuracy and specificity of calls made by both platforms for ONT pipelines. For each, calls made by all but one pipeline (or both technologies) were considered true positives, while calls made by only a single pipeline (or technology) were considered false positives, thus the ROC AUCs cannot be directly compared between groups.

To ensure the approach employed was robust to changes in the virus and sequencing methodologies over time, a more recent set of samples (collected from 04/14/2021 to 04/02/2022) were analyzed as well after having settled on our normalization approach. To determine the extent of agreement and disagreement across pipelines, variants calling results were compared and plotted as upset plots (**Figure 4).**First, results without any filtering applied were considered, **Figure 4A-D.**Here only variants with an AF of at least 15% were considered and the results had both SPDI and Parsimony normalizations applied. While Illumina data showed strong agreement, even prior to filtering for both SNPs and InDels (**Figure 4A+B**, respectively), for ONT data, for both SNPs and InDels (**Figure 4C+D** respectively), the results were dominated by pipeline unique calls, as evidenced by the higher bars to the left of the graph where, below the bars, there is a single filled dot associated with a single pipeline. Next, we applied the filtering outlined above, **Figure 4E-H**. For both Illumina and ONT SNPs (**Figure 4E** and **Figure 4G** respectively), the filtering increased the concordance in variant calls, though this was more pronounced for the ONT results. Thus, the majority of pipeline discrepancies can be attributed to calls with AF < 50%, calls at locations with DP < 100 or an ALTDP < 50, calls from samples with poor reference genome coverage (<50%), calls in homopolymer regions, or from single-end data. While the approach did improve the agreement in InDel calls for both Illumina and ONT (**Figure 4F** and **Figure 4H** respectively), the extent of agreement was not as strong as seen for SNPs. Notably, the total number of InDel calls was greatly reduced, suggesting that the majority of InDel calls may be artifactual or otherwise poorly supported. The initial test set also showed good agreement in variant calls postfiltering (**Supplement Figure 2**), however the improvement in InDel calls was less pronounced suggesting that the differences in Dataset1 may have been due to other issues. Additionally, agreement in both SNP and InDel calls across the length of the genome was plotted as a heatmap (**Supplemental Figure 3**) and no significant association with any portion of the genome was observed.

**Figure 4.**
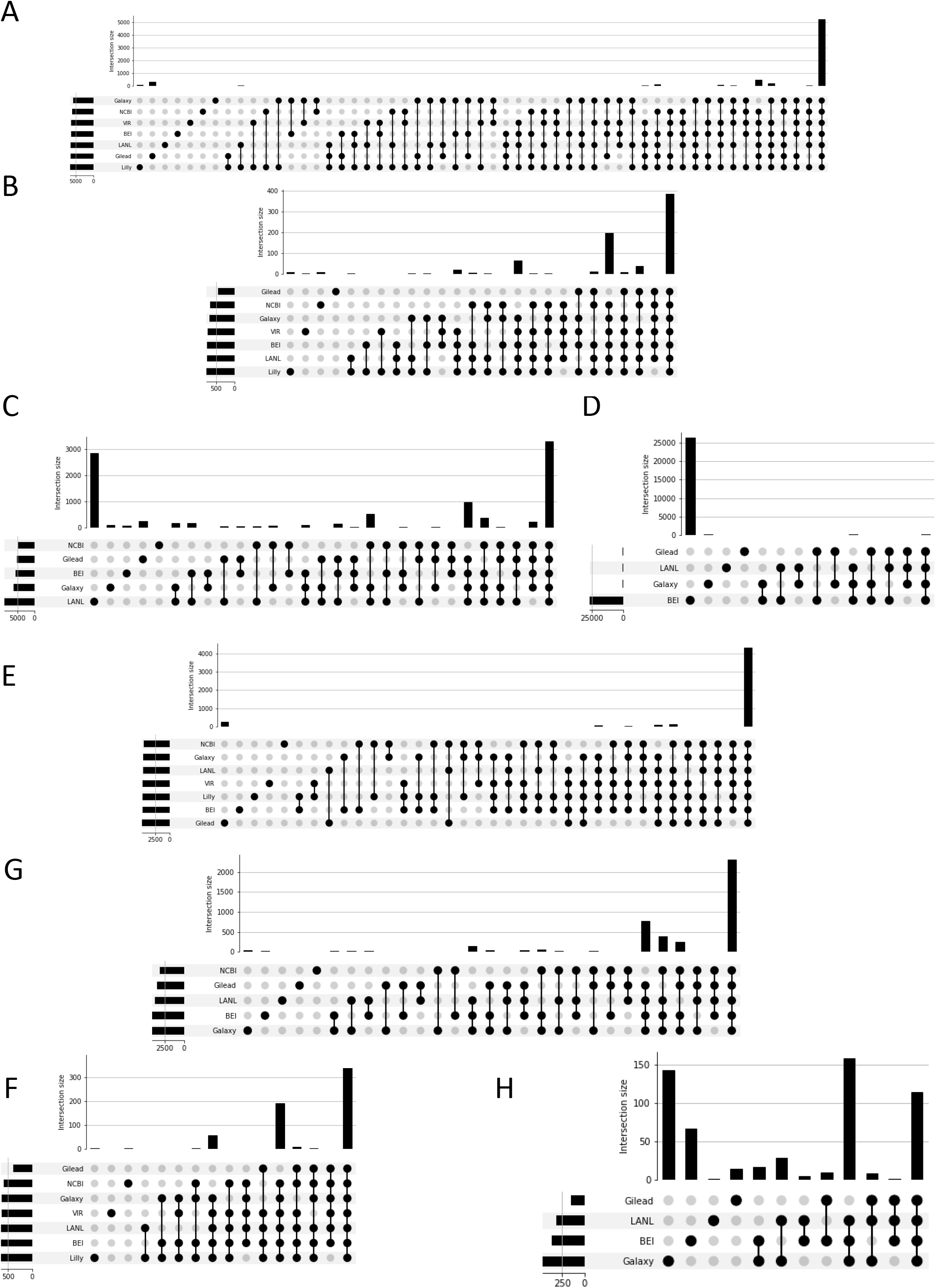
Agreement across pipelines with and without recommended parameters. A+B+C+D) Agreement across pipelines without recommended parameters. E+F+G+H) Agreement across pipelines with recommended parameters. A+E) Agreement on Illumina SNP calls. B+F) Agreement on Illumina InDel calls. C+G) Agreement on Oxford Nanopore (ONT) SNP calls. D+H) Agreement on ONT InDel Calls. For each figure, the bars indicate the number of variants called by the groups indicated by filled circles below, across the whole dataset. The large number of workflow unique calls for LANL and BEI in C) and D) respectively are attributable to calls with low read support (DP<100 or ALTDP<50), of low frequency (<50%), in homopolymer regions, or in samples with poor reference coverage, as indicated by the reduction after filtering, G) and H) respectively.

Similarly, to assess the extent of agreement and disagreement across sequencing technologies, stacked bar plots were generated (**Figure 5**), with samples being considered only if both Illumina and ONT data passed the filtering criteria described above. Similar to what was seen with the cross-pipeline comparison, the results for both SNPs and InDels without filtering (**Figure 5A** and **Figure 5B** respectively) were markedly improved for both SNPs and InDels with the application of our filters (**Figure 5C** and **Figure 5D** respectively). Again, similar to what was seen with the cross-pipeline comparison, the initial dataset showed similar agreement for SNPs and more modest agreement for InDel calls post filtering (**Supplemental Figure 4**), again suggesting that the discrepancies observed with the initial data set are likely due to issues in sequencing early during the pandemic.

**Figure 5.**
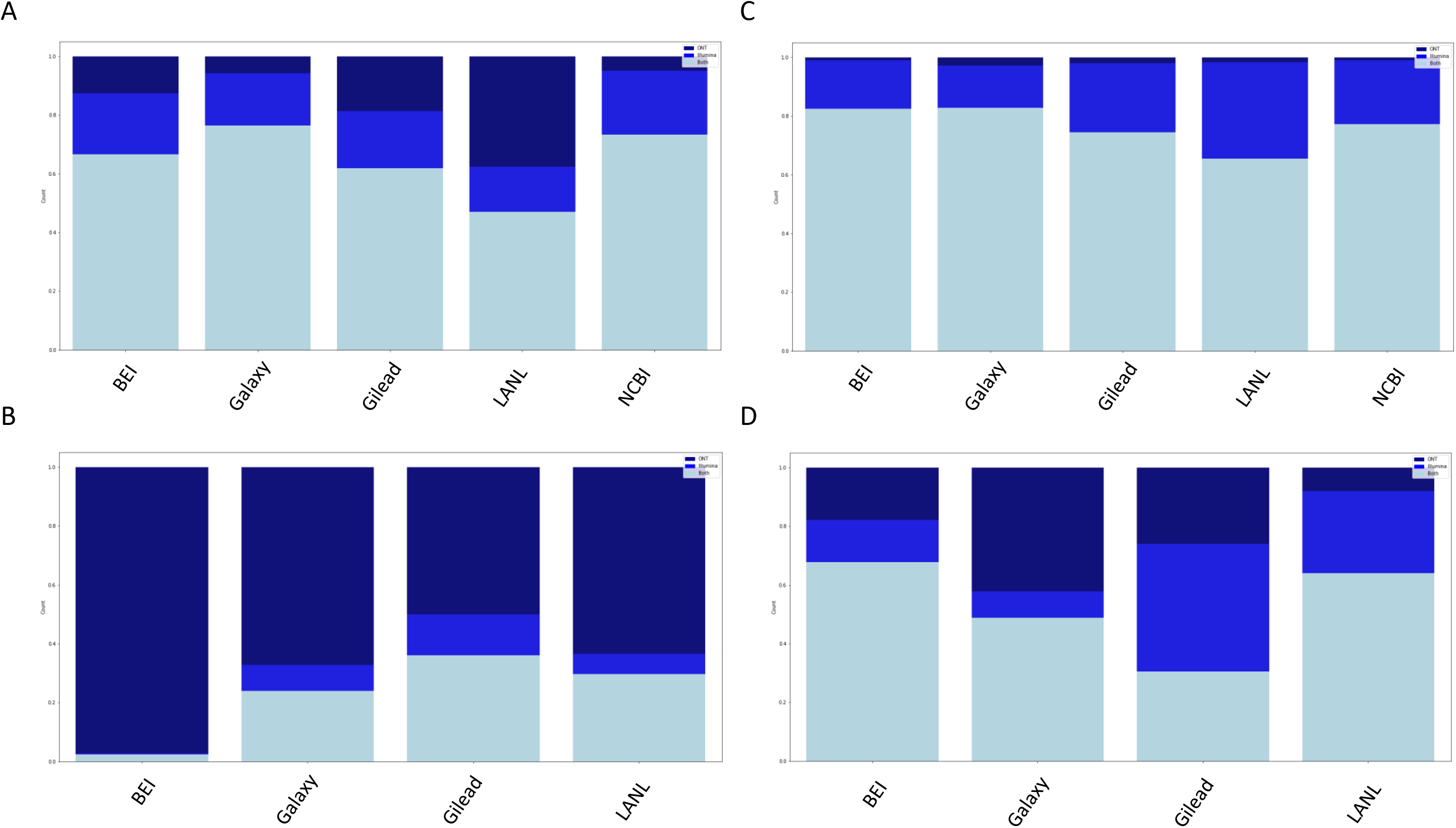
Agreement across platforms with and without recommended parameters. A+B) Agreement between platforms without recommended parameters. C+D) Agreement between platforms with recommended parameters. A+C) Agreement between platforms on SNP calls. B+D) Agreement between platforms on InDel calls. For each figure, only those sample for which both Illumina and ONT platform data had at least one variant call that passed all the filters was considered. The total height is normalized to the total number of calls made by each pipeline, with the light blue portion indicating calls made on both platforms for a given sample, the medium blue indicating calls made only for the Illumina data, and the dark blue indicating calls made only for the ONT data.

In summary, the filtering we identified (summarized in **Figure 6**), provides robust agreement across diverse variant calling pipelines for both Illumina and ONT platform data and supports robust agreement between these technologies. This in turn should support consistent reporting of consensus sequences for downstream analyses, and the fair comparison of results generated by many different sequencing groups.

**Figure 6.**
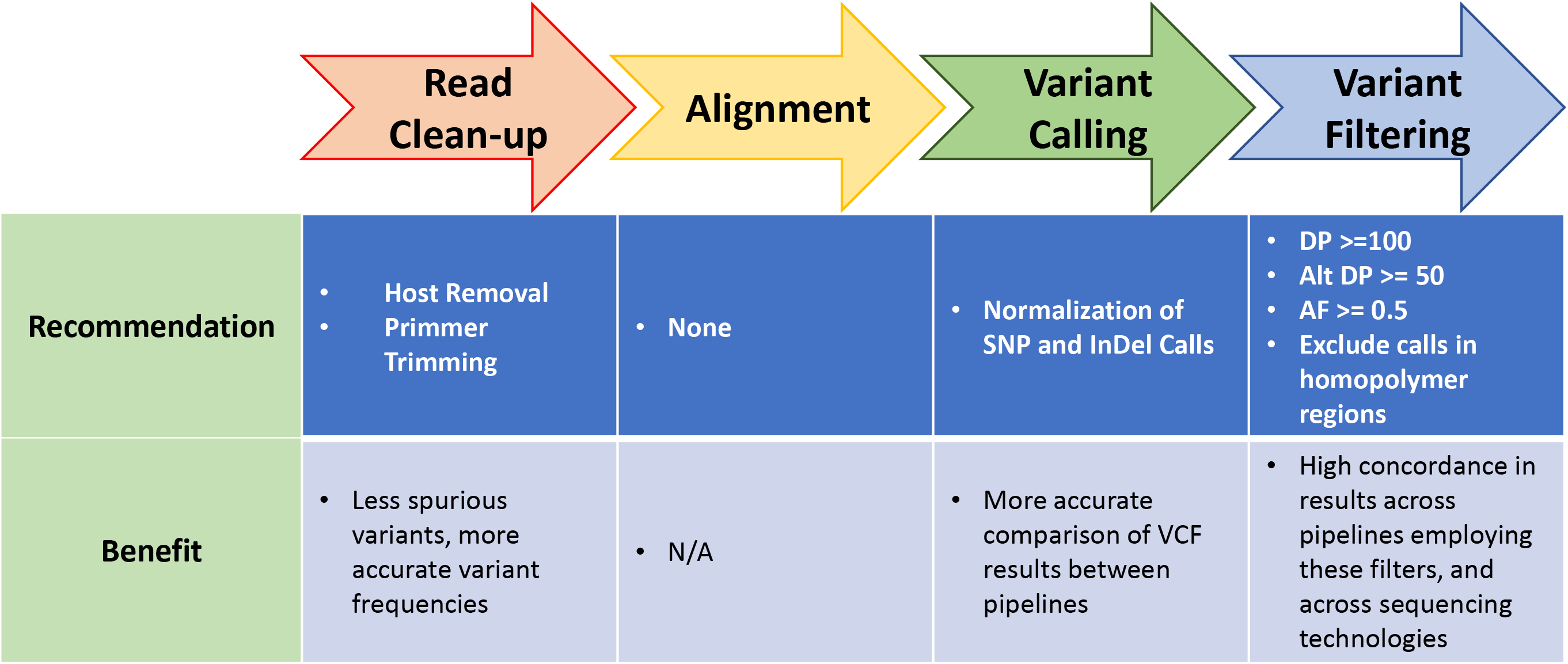
Variant Calling Pipeline Recommendations. Recommendations for each step in a variant calling pipeline, from read cleanup to variant filtering, are illustrated. Additionally, the benefit of implementing the recommendations at each step are noted.

## Discussion

In the current pandemic, at least at the beginning, the majority of the effort was focused on analyzing consensus genomes. However, recent cases of possible recombinants, co-infection, contamination, and monitoring for drug resistance to antivirals has made clear the importance of working beyond consensus genome.^14^ Consensus genomes reduce the complexity and size of the data, but this reduction also removes information that may be pertinent to determine the quality (contamination, bioinformatic workflow, etc.) of data and increases the difficulty of extracting relevant information (co-infection, minority variants, recombination, etc.) from the data. Coinfections and contamination, which can come in many forms and can happen during sample collection, laboratory preparation, sequencing, or bioinformatics processing,^15–18^ if not detected during consensus genome generation, can get propagated as biological signal. In absence of corresponding raw data, such issues cannot be easily tracked and corrected. Still, as consensus genomes are the starting material for many common analyses, we aimed here to identify factors which influence the accuracy and consistency of variant calling results.

A key feature of the study presented here is the variety of sample types and bioinformatics analytical pipelines. Every sequencing methodology has some degree of implicit bias and error profile toward genomic regions, such as regions containing homopolymer repeats or near the ends of linear genomes. Biases can also be introduced by upstream sample handling procedures, including different library preparation and DNA/RNA extraction techniques, as well as downstream analysis methods, ie. algorithms used by different bioinformatics tools. Next Generation Sequencing is an increasingly valuable technology that provides a great deal of analytical insight, but it comprises a series of different methods and few, if any, are objectively superior or all encompassing. To gain the clearest representation of any given genome, more diverse sequencing methods and more diverse analytical pipelines should be employed. Though an evaluation of library preparation methods is outside the scope of this work, that remains a significant point of interest in determining the validity of any detected variant through sequencing methods. With this study including seven different groups reviewing the same sequencing data, themselves derived from a much greater breadth of institutions and methods, we are able to show the degree of concordance between these methods and technologies and infer meaning. Illumina is known for its accuracy in short-read (75-300 base pairs) sequencing^19^. A high accuracy rate is ideal for determining small nucleotide variations relative to another genome sequence. However, short read data is challenging to interpret in convoluted sequence regions, which are prevalent in many genomes^19,20^. In the event a region of a genome is difficult to deconvolute over a range longer than that of a single read, a bioinformatics alignment program will struggle to place this read in the correct location. Oxford Nanopore Technologies (ONT) produces reads of variable lengths that are typically much longer than their Illumina counterparts. These reads are ideal for spanning the previously described convoluted regions of a genome, however they have a higher error rate^19,20^. From a variant calling perspective, reads are more likely to be accurately placed but more likely to have infrequent variant artifacts found at a given position.^21^ While Illumina technology has been most frequently used in next generation sequencing and many variant callers have been designed and optimized to specifically handle Illumina reads, ONT sequencing is comparatively new and actively being improved, including updates that reduce the error rate, with lower sequencing cost (https://nanoporetech.com/about-us/continuous-development-and-improvement). Long and short read technologies are being increasingly linked in hybrid assembly software to benefit from the strengths of each technology while minimizing the weaknesses. In a similar vein, a variant calling effort that compares those called on Illumina reads and those called on ONT reads would find high confidence in any variants called on both platforms for the same samples.

The ACTIV-TRACE group employs different alignment and variant calling pipelines from each contributor. Variant calling tools tend to be built for specific purposes, such as human genomics, microbial genomics, or metagenomics. Alignment tools are less deliberately biased toward organism type but employ different algorithms to handle specific read types.^21^ Within both alignment and variant calling algorithms, there are explicit and implicit filters related to match score, mismatch penalty, strand-bias handling, depth and frequency requirements, and primary alignments that further differentiate even identical programs between pipelines. The benefits of several pipelines are not limited to variant calling and alignment. Standard pre-processing steps of adapter or base quality trimming, de-hosting or taxonomic identity binning, and read deduplication or complexity requirements add further nuance to results comparison. The ACTIV-TRACE group’s variety in analytical methods provides the opportunity to examine how differently they perform in the context of SARS-CoV-2.

While the work here is focused on variant calling, the major phylogenetic approaches employed extensively during the current pandemic rely on consensus sequences as input data.^22,23^ As consensus sequences can be viewed as a condensed representation of variant calling data, it is worth noting some of the implications of the current work on that data type. Firstly, while the various pipelines employed here achieved excellent consensus, there were still differences between each pipeline (e.g. the number of False positives found by each pipeline after filtering, varies from less than 3% of remaining calls to approximately 27%). Thus, it is important that information on how sequencing was conducted, and how the subsequent bioinformatic analyses were conducted be deposited along with the consensus sequences so that analysis using them can take the information into account in determining if the sequences are truly comparable. Further, primer trimming can have a notable effect on variant calling results, as improperly trimmed reads will result in incorrect variants that reflect the primer sequence instead this information should also be included with consensus sequence submission. Additionally, given that the discrepancies between pipelines is greater at lower allele frequencies, and that consensus sequences cannot capture the detail found in VCF’s beyond the use of IUPAC ambiguity codes, it is recommended that a relatively high allele frequency be chosen for inclusion into a consensus sequence.^24^ While this recommendation is expected to increase the comparability of consensus sequences generated across pipeline, it may reduce the useful of consensus sequences for drug resistance monitoring. An allele frequency threshold of 50% will ensure that mutations included in consensus sequences are of high confidence and reduce the need for us of IUPAC ambiguity codes, which themselves present challenges for many downstream bioinformatic applications. Similarly, as consistency across pipelines is sensitive to overall and alternate allele depth, requiring a minimum depth of 100 and alternate allele depth of 50 should ensure that those mutations that make it into a consensus sequence are of high confidence. For cases in which a minimum depth is not met, until variant calling software enables the usage of IUPAC ambiguity codes in reference sequences, these ambiguous locations should be reported as “N.” Together, these recommendations would help ensure that the results represented in consensus sequences are of high confidence, i.e., are likely replicable across different pipelines and sequencing technologies, and thus are appropriate to compare to each other during downstream analyses.

Of note, having the same dataset being analyzed by multiple groups offered the possibility for each group to prioritize the investigation/visual assessment of potentially spurious variant calls (based on the comparison with the results of the other groups) and tune their pipeline parameters accordingly, with the acknowledged limit that a variant being called by majority of the pipelines does not necessarily mean it is real (example sample contamination or library preparation artefact).

Pharmaceutical and biotechnology companies are required to comply with Good Practice (GxP) guidelines and obtain system validation, when developing computer software to ensure that medical devices, drugs and other life science products are safe and effective (https://www.fda.gov/regulatory-information/search-fda-guidance-documents/part-11-electronic-records-electronic-signatures-scope-and-application). At present, it is unclear to what level of computer system validation is required by the FDA for NGS analysis pipelines, and the FDA performs their own independent analysis and compares to submitted analysis from sponsor. Given the numerous sample preparation methods and sequencing technologies as well as the various analysis pipelines (many of which are proprietary), this presents challenges for regulatory review of NGS data.^25^ In the future, the FDA anticipates the development of standardized NGS analysis pipelines that will provide a reproducible data analysis sufficient for generating consistent and robust results from NGS data.^25^ Based on the analysis described here within, standardized NGS analysis pipelines have not been achieved given the lack of agreement in the analysis of publicly available NGS datasets. Further collaborative efforts like ACTIVE TRACE’S work here are needed to achieve standardization of pipelines and reproducibility of results.

## Online Methods

### Common Features

RefSeq^26^ accession sequence NC_045512.2 was used as reference genome for alignment. A description of the data analyzed in the original and recent datasets can be found in **Supplemental Table 2** and **Supplemental Table 3**, respectively. Briefly, Dataset 1 consists of 413 sequence records, representing 155 samples, while the recent set consisted of 419 sequence records, representing 210 samples. Dataset 1 consists mostly of pre-alpha lineages and some alpha samples, while Dataset 2 has a mix of VOCs (alpha, delta, omicron and some other lineages). Both sets are constituted by paired Illumina and ONT samples. The records were processed by each of seven pipelines described below. The results were then combined, and SNPs were normalized using the SPDI algorithm.^13^ Subsequently, the InDel results are normalized using the Parsimony script, described below, and the additional analyses and figures were generated via python scripts, all available here (https://github.com/ncbi/ACTIVTRACEvariants).

### Parsimony Script

Following SPDI-normalization, the parsimony script is applied. After pulling the SPDI-processed data into a data table, the first step is to sort data according to analytical group, sample accession, and position. Then, it must resolve adjacent indels into singular records. Where InDels use a hyphen (-) to represent the reference or alternate allele, we search for any adjacent rows with identical group and accession values and the positions are consecutive. For each record that satisfies these requirements for either the next or previous record, they are grouped together and concatenated according to the Group and Acc values, with any number of repeated hyphens being replaced with a singular hyphen to match SPDI formatting. This only saves the depth and alternate allele values for the first record in the set of each consecutive InDel. In practice, we have not found this disruptive to our analysis, but this may present an issue when this occurs closer to the limit of detection or when precise depth is significant.

Next, we must address the nucleotide context issue. This combined data table is looped over, row by row, capturing the assorted metadata in each row. These fields will not be altered. We are only concerned with InDels that don’t conform to our simple formatting. First, we check for the simple cases that either the reference or alternate allele completely contains the other, such as AAA4313AAAA. In this case, we remove the shorter of ref or alt from the other and replace the shorter field with a hyphen (-) and write to file, transforming AAA4313AAAA to A4313-.

This leaves the more complex case of a single record containing both an InDel and a SNP. These records have neither a reference nor alternate allele that completely contains the counterpart, so we cannot simply remove the identity of one from the other, as we had before. Instead, we align the reference allele and the alternate allele for each qualifying row using biopython’s^27^ pairwise2 module as align.globalms with settings match=2, mismatch=-0.5, gapopen=-1, gap extension=0.1 and allow only the top alignment. These settings were determined to be sufficient for the data presented in this study but have not been optimized to other datasets. For each alignment, we determine where a deletions may be found and save InDel in the ‘type’ field of the metadata variable. For all other nonmatching bases in the alignment, we save SNP to the ‘type’ field of the metadata. In both cases, the position of the variant is incremented by the number of preceding matching bases to correct for where the variant occurs. These explicit and separated InDel and SNPs are written as discrete records to file. In all remaining cases, both the reference and alternate allele values match with at least the starting nucleotide of the record. We remove all matching alternate alleles from the reference alleles that haven’t been otherwise addressed from the beginning of the reference and increment the position by the length of the removed bases. This will leave only a deletion in all observed cases. These are written to file with a hyphen in place of the alternate allele.

### Studying the effects of primer trimming on variant calls and apparent allele frequencies of called variants

83 high-quality ARTICv3-amplified samples were selected from the ACTIV TRACE Illumina paired-end sample collection. These samples were analyzed twice with the Galaxy pipeline, once with and once without primer trimming. The position of each resulting variant call (2696 calls total with, 2637 without trimming) was compared to the known primer binding sites of the ARTICv3 primer scheme to classify calls as inside (421 calls with, 360 calls without primer trimming) and outside of primer binding sites. For variants called with and without primer trimming, the observed variant allele-frequency with primer trimming was plotted against the same metric without primer trimming. For variants called only with primer trimming a value of zero was used as a substitute for the unobserved variant allele-frequency without primer trimming.

### Calculation of Receiver Operating Characteristic (ROC) plots based on concordance across pipelines

For each participating group, all variant calls made for all samples analyzed with the group’s pipeline were classified as either concordant or discordant based on whether that same variant had or had not been called for the same sample by the majority of groups (>= 6 pipelines) that had analyzed that sample. For the purpose of generating ROC-like plots, concordant and discordant calls were treated as true-positive and false-positive calls respectively. The true- and false-positive lists of each group were then filtered independently with increasing thresholds on two key variant call metrics: the number of sequencing reads supporting the variant allele (alternate allele read depth, AltDP) and the fraction of all sequencing reads at the variant site that support the alternate allele (alternate allele-frequency, AF). Increasing thresholds of each of the two metrics lower the number of true- and false-positive calls, but to different extents. For plotting, the numbers of retained true- and false-positive calls at each threshold were normalized to the numbers of unfiltered true- and false-positive calls of the respective group, thus the ROC AUCs cannot be directly compared between groups.

### Calculation of Receiver Operating Characteristic (ROC) plots based on cross-platform agreement

For samples for which both Illumina and ONT sequencing data was available, alternative plots could be generated as follows.

For each participating group, all variant calls made for all samples analyzed with the group’s pipeline based on the data for one of the two sequencing platforms were classified as concordant or discordant based on whether that same variant had or had not been called for the same sample on the other platform either by any participating group, or not. For plotting, the true- and false-positive lists of each group resulting from this alternate classification were used to create a threshold and subsequently normalized as described above.

### BEI Resources

SARS-CoV-2 reads were retrieved from NCB’s^28^ Sequence Read Archive (SRA) directly using the fastq-dump v2.11.1 utility (https://github.com/ncbi/sra-tools). These reads were then trimmed and filtered to remove adapter sequences and low-quality reads, using *fastp*^29^ v0.23.2 for Illumina reads and *NanoFilt*^30^ v2.8.0 for Oxford Nanopore reads. Settings for *fastp* were left at default, while *NanoFilt* was set for a minimum average quality of 10 and minimum length of 150, while trimming the first 30 bases of each read. Following this, both sets of reads were taxonomically binned using ATCC’s *bin_reads* function, which relies on *kraken2*^31^ v 2.1.2 to identify the nearest taxonomy of any given read. Kraken2 classification was run with default settings and its bacterial_viral_db database. Kraken2 also contains an extract reads function that was employed using NCBI: taxID6940099, which corresponds to SARS-Coronavirus and all sub-taxa.

All Illumina reads that have been processed as outlined above were entered into an identical pipeline for variant calling analysis. This pipeline comprises four steps: read mapping to NC_045512.2 (Wuhan-Hu-1), local realignment, variant calling, and normalization. Prior to alignment, paired-Illumina fastqs need another step of processing. Because kraken2’s taxonomic binning does not guarantee that both forward and reverse fastqs will have the same resulting reads, *seqkit common*^32^ v 2.1.0 is used to ensure that only paired reads are further analyzed. Alignment was performed with *bwa mem*^33^ v 0.7.17-r1188 using default parameters. Local realignment begins with *bcftools mpileup*^39^ v1.12 to produce a per-base pileup to stage the realignment. This uses default settings, except for the max-depth which was set to 8000 to reduce the chance of missing minority variants due to excessive depth at a position. Next, *bcftools call* and *bcftools filter* are used to capture multiallelic variants with a variant call quality above 30. This initial vcf is passed to *GATK*^35^ v4.2.2.0 to apply base quality score recalibration within the context of all bases at each position. Finally, realignment is achieved using the *lofreq viterbi*^36^ tool v2.1.5, using default parameters. Variant calling is performed using *lofreq* v2.1.5 in four stages: *indelqual* with “--dindel” parameter and *alnqual* to capture indel variants, *call* with --call-indels and -C 50 to capture all variants above a depth of 50x, *and filter* with -v 50 and -a 0.15 to further filter for minimum depth of 50x and alternate allele frequency of 15%.

Oxford Nanopore long reads have a few fundamental differences that affect what off-the-shelf tools can be used for their processing. Namely, underlying algorithms in *fastp, GATK*, and *bwa mem* are not designed for the increased length and error profile of these reads. Consequently, medaka_haploid_variant^37^ v1.5.0 is used to perform the initial alignment and its BAM file is passed to lofreq in the same manner as in the Illumina dataset. Minimum depth and alternate frequency filters are applied afterward and the previously described step of taxonomic binning is identical to the Illumina dataset.

### Galaxy Project

Illumina short-read (paired-end subset of the data only) and Oxford Nanopore long-read data was downloaded in fastq.gz format from the FTP server of the European Bioinformatics Institute (EMBL-EBI) at ftp.sra.ebi.ac.uk. Variant calling was then performed with platform-specific Galaxy workflows, which have previously been described^38^ and are publicly available from Dockstore^39^ and the WorkflowHub^40^ against NCBI Reference Sequence NC_045512.2. No attempt was made to analyze single-end Illumina data.

A brief overview of the analysis of data from both platforms is provided below.□

#### Variant calling from paired-end Illumina short-read data

We used fastp (version 0.23.2) for read trimming and quality control^29^, aligned reads with bwa-mem (version 0.7.17)^33^, filtered for reads with fully mapped read pairs (with samtools, version 1.9^34^), re-aligned reads using the lofreq viterbi command from the lofreq package (version 2.1.5 used here and in all subsequent steps using lofreq)^36^ and calculated indel quality scores with lofreq indelqual. We then attempted to trim amplification primers from the aligned reads using ivar trim (version 1.3.1)^41^ assuming the ARTIC v3 primer scheme^42^ had been used in amplification of all samples (which is true for the majority of the samples analyzed, but not all of them; we continued with untrimmed data for those other samples). The remaining, trimmed alignments served as the input for variant calling with lofreq, and variant calls down to an allele-frequency of 0.05 were reported if confirmed by at least 10 reads.

#### Variant calling from Oxford Nanopore long-read data

We used fastp (version 0.23.2) for quality control^29^ and read length filtering. For samples that we detected to be amplified with the ARTIC v3 primer scheme we filtered for read sizes between 300 and 650 bases, for other samples we allowed read sizes between 300 and 3,000 bases. Retained reads were aligned with minimap2 (version 2.17)^33^, and successfully mapped reads left-aligned with BamLeftAlign from the freebayes package (version 1.3.1)^43^, after which we attempted to trim primers from the reads with ivar trim (version 1.3.1), again assuming the ARTIC v3 primer scheme and using untrimmed data where that assumption failed. The data was then analyzed with the medaka consensus tool (https://github.com/nanoporetech/medaka, version 1.0.3)^37^ and variants extracted with medaka variant tool (version 1.3.2) and postprocessed with medaka tools annotate (integrated into the medaka variant tool’s Galaxy wrapper; https://toolshed.g2.bx.psu.edu/repository?repository_id=a25f9bf8a7d98ae4&changeset_revision=0f5f4a208660). Variant calls down to an allele-frequency of 0.05 were reported if confirmed by at least 10 reads and called, in the case of SNVs only, with a QUAL score of at least 10.

### Gilead Sciences

Illumina short-reads and Oxford Nanopore long-read data were downloaded from Sequence Read Archive (SRA) using sratoolkit (https://github.com/ncbi/sra-tools#the-sra-toolkit, v 2.8.1).

#### Illumina□

Fastq files were aligned to hg38 reference using BWA v0.7.15^33^ to exclude human RNA transcripts and to isolate viral reads for further processing. Next, reads were trimmed using Trimmomatic v0.3 6^44^ for low quality (sliding window 4 bp, avg phred 15) and short reads (<50 base pairs) were filtered out. Paired end reads that overlap were merged using NGmerge v0.3^45^ software creating a single-end fastq file containing merged reads and any single end reads that do not overlap. Reads are then aligned to the Wuhan-Hu-1 reference (NC_045512) using SMALT v0.7.6 aligner (https://www.sanger.ac.uk/tool/smalt-0/). If amplification primer information was available, trimmed the base pairs from reads that overlap with primers. Tabulate nucleotide variants and indels per genome position (NC_045512), excluding any variants with average phred score less than 20 and read depth less than 50 as well as any frameshift indels.

#### ONT

Fastq files were aligned to hg38 reference using minimap2 v2.17^46^ to exclude human RNA transcripts and to isolate viral reads for further processing. Reads are then aligned to the Wuhan-Hu-1 reference (NC_045512) using minimap2 v2.17.^46^ If amplification primer information was available, trimmed the base pairs from reads that overlap with primers. Tabulate nucleotide variants and indels per genome position (NC_045512), excluding any variants with average phred score less than 10, forward strand ratio < 0.1 or > 0.9, and read depth less than 50 as well as any frameshift indels.

### LANL

Illumina short-read and Oxford Nanopore long-read data were downloaded from Sequence Read Archive (SRA) using sratoolkit (https://github.com/ncbi/sra-tools#the-sra-toolkit, v 2.9.2). Quality control, read mapping, variant calling, and consensus genome generation were then performed with EDGE-COVID19 (EC-19) workflows (http://edge-covid19.edgebioinformatics.org/). Detailed description of the EC-19 workflows has also been previously described^47^ (https://edge-covid19.edgebioinformatics.org/docs/EDGE_COVID-19_guide.pdf). Briefly, the EC-19 workflow employs many commonly used tools such as FaQCs (v2.09) for quality control, Minimap2 (v2.17, default for ONT) or BWA mem (v0.7.12, default for Illumina) for mapping reads to a SARS-CoV-2 reference genome (NC_045512.2 without the 33nt poly-A tail in the 3’ is used as default), an algorithm based on ARTIC pipeline (https://github.com/artic-network/fieldbioinformatics) for trimming primers if an amplicon-based method (e.g. ARTIC, SWIFT) was used for sequencing, generating consensus genomes, and variant calling based on samtools mpileup (v1.9) wrapped into a custom script (https://gitlab.com/chienchi/reference-based_assembly). EC-19 then accounts for strand biasness using both fisher score and Strand Odds ratio (https://gatk.broadinstitute.org/hc/en-us/articles/360036464972-AS-StrandOddsRatio) and reports SNVs if the Allele Frequency (AF) > 0.2 and have minimum Depth of Coverage (DC) of five5. Likewise, InDels are reported if DC > 5 and then platform specific thresholds are implemented for AF as AF>0.5 is required for Illumina and AF> 0.6 for ONT data in order to account for the higher error rates with this technology, and >0.8 within homopolymer sequences.^48,49^

### Lilly

#### Illumina

The paired end raw sequencing data were trimmed in two successive rounds using cutadapt^50^ version 2.5 with the following parameters:□]

pe1_parms:□ --quality-base=33 -a ‘G{150}’-A ‘G{150}’:□□□

pe2_parms: -a AGATCGGAAGAGCACACGTCTGAACTCCAGTCAC -A AGATCGGAAGAGCGTCGTGTAGGGAAAGAGTGTAGATCTCGGTGGTCGCCGTATCATT : --quality-base=33 -q 20 --n 2 -trim-n -m 20 --max-n=.2 : -a ‘{150}’ -A ‘{150}’ -a ‘T{150}’ -A ‘T{150}’ : -g ‘{150}’ -G ‘{150}’ -g ‘T{150}’ -G ‘T{150}’: -g AAGCAGTGGTATCAACGCAGAG -G AAGCAGTGGTATCAACGCAGAGI3 : -g AAGCAGTGGTATCAACGCAGAGTAC -G AAGCAGTGGTATCAACGCAGAGTAC□□

Reads were aligned to the hg19_human_trxome_and_coronavirus.fa reference genome (Thermo Fisher, which included MT019532.1 BetaCoV/Wuhan/IPBCAMS-WH-04/2019) using bwa-mem^33^ version 0.7.12 with default parameters. Variants were called using FreeBayes^43^ version 1.3.1 with the following parameters: -F 0 -p 1 -K -C 0 -n 5 -w --min-alternate-count 0 --min-alternate-fraction 0.

Variants were reported if there were >= 5 reads supporting the variant; if□ >= 10% of reads supporting the variant were derived from the minor strand; and if the allele frequency was >=15%.□

### NCBI

The NCBI SARS-CoV-2 variant calling pipeline can be found at https://github.com/ncbi/sars2variantcalling. The pipeline, as used for the analyses presented here, is briefly described below.

#### Illumina

Illumina short reads were downloaded from SRA database using fastq-dump of sratoolkit (https://github.com/ncbi/sra-tools#the-sra-toolkit, version 2.11.0), then trimmed using trimmomatic (version 0.39)^44^. The reads were aligned using Hisat2 (version 2.2.1)^51^, then left-aligned using GATK LeftAlignIndels (version 4.2.4.1)^52^. GATK HaplotypeCaller (version 4.2.4.1) with the options “minimummapping-quality 10”^52^ was used for generating variant VCFs with NCBI Reference Sequence NC_045512.2 as the reference. Calls with QUAL value smaller than 100, alternate allele counts lower than 10, FS value smaller than 60, SOR value smaller than 4, QD value equal to or greater than 2, ReadPosRankSum value equal to or greater than −4, allele frequency lower than 0.15, and reference genome positions beyond 29850 were excluded.□

#### ONT

Nanopore reads were downloaded from SRA database using fastq-dump of sratoolkit (https://github.com/ncbi/sra-tools#the-sra-toolkit, version 2.11.0), then trimmed using NanoFilt^30^ (version 2.8.0) with the options “q 10” and “headcrop 40”^30^. Two rounds of medaka (version 1.3.2, https://github.com/nanoporetech/medaka) were applied to generate consensus assembly. Consensus to reference (NC_045512.2) alignment and initial variant calls were generated with MUMmer (version 4.0.0rc1, https://github.com/mummer4/mummer). InDels and SNPs within 10bps of an InDel were excluded using bcftools (version 1.11)^34^, and snp clusters (2 or more SNPs within 10 bps) were filtered using vcftools (version 0.1.12b),^24^□□

□

### VIR□□□□□

#### Illumina□

The pipeline was written and optimized for another library preparation and the primer removal step and pipeline parameters were not optimized for the samples analyzed in this study.

Illumina short reads□were downloaded from SRA□database□using□fastq-dump of□sratoolkit□ (https://github.com/ncbi/sra-tools#the-sra-toolkit, version□2.9.1) [parameters: –*split-files* for paired end reads].□

The library preparation consisted of a mix of SE reads, and PE reads (with random fragmentation or amplicons fragment). The read length ranged from 300 to 500bp for SE, and 50 to 300bp for PE reads. As the samples were prepared with different library preparations, not always retrievable from the metadata, we used a conservative approach to trim the 31bp at the beginning of all reads that were 150bp or longer. 31bp corresponds to the maximum primer length in the ARTICV3 kit. Reads were trimmed with trimmomatic (version 0.39) [parameters: HEADCROP:31 MINLEN:35; PE -validatePairs for paired end; SE for single end].^44^

The alignment was performed with bwa-mem (quay.io/biocontainers/bwa:0.7.17--hed695b0_7) [parameters: -M].^53^ The variant calling was performed with lofreq (quay.io/biocontainers/lofreq:2.1.5--py36ha518a1e_1)^36^ in multiple consecutive steps: lofreq viterbi; lofreq indelqual [parameters: --dindel]; lofreq call-parallel [parameters: --no-default-filter --call-indels –min-bq 6 –min-alt-bq 6 –min-mq 1 –sig 1]; lofreq filter [parameters: --no-defaults –af-min 0.01 –cov-min 15 –sb-mte fdr-sb–alpha 0.05 –sb-incl-indels; note: --sb-alpha parameter was set to 0 for samples prepared with amplicon libraries to prevent filtering variant due to strand bias, which can occur with amplicon library preparation]. For each sample, SNVs present at AF>0.5 were substituted in the reference genome with bcftools consensus (quay.io/biocontainers/bcftools:1.10.2-hd2cd319_0)^34^ and the alignment and variant calling steps were run a second time on the “new” reference genome (in order to rescue reads that were potentially mis-aligned/un-aligned due to too many mutations located in the near vicinity of each other). Of note, the final variant calling is done with regards to the initial NC_045512.2 reference genome nomenclature. SNVs and indels were reported if they were present with AF>0.15. Minimum read depth was set at a low threshold (15 reads) for the purpose of this analysis. Variants flagged as potentially spurious due to their location on the reads (i.e. consistently located at the same position in the read for the alternative allele, but not for the reference allele) were further filtered if they were present in more than 2 samples and in all samples at low AF (<0.5).

#### ONTI3

At the time of analysis, VIR did not have an existing pipeline to assess ONT data.

## Acknowledgements

This work was supported in part by the National Center for Biotechnology Information of the National Library of Medicine (NLM), National Institutes of Health. This work was supported in part by the European Union’s Horizon 2020 and Horizon Europe research and innovation programs under grant agreements No 871075 (ELIXIR-CONVERGE) and 101046203 (BY-COVID). This project was funded in part with Federal funds from the National Institute of Allergy and Infectious Diseases, National Institutes of Health, Department of Health and Human Services, under Contract No. HHSN272201600013C, managed by ATCC. This work was supported in part by the Los Alamos National Laboratory’s Laboratory-Directed Research and Development program [20200732ER and 20210767DI] and under IAA project RRJJ00 with the Centers for Disease Control and Prevention. We would like to acknowledge Roberto Spreafico, Leah Soriaga, Li Yin and Amin Momin for their contribution in the VIR variant calling pipeline, and Lisa Purcell for providing guidance and advice to this group. We would also like to acknowledge John Calley for his creation of the Lilly variant calling pipeline. The work would not be possible without the NIH ACTIV TRACE initiative and the support of the Deep Sequencing Analysis Subgroup and the HHS Office of the Assistant Secretary for Preparedness and Response (ASPR) Countermeasures Acceleration Group (CAG).

## Conflict of Interest

RM, JL, and ESM are employees of, and stockholders in, Gilead Sciences, Inc. JdI is an employee of, and hold stock or stock options in, Vir Biotechnology, Inc. PE is an employee of, and holds stock or stock options in, Eli Lilly and Company. The remaining authors declare no conflict of interest.

## Author Contributions

RC, DAY, WM, MS, RM, RB, JRB, PSGC, CAC, JdI, PE, TLR, PL, CL, SR, ESM, CX, and KDP all contributed to editing the manuscript. DAY, WM, MS, RM, RB, JdI, BH, PE, YJ, JG, KSK, AK, JL, PL, CL, SR, CX, VZ all contributed to the development of one of the workflows and/or processed data used in this study by one of the workflows. RC, DAY, WM, MS, RM, PSGC, JdI, PE, JL, PL, CL, CX all conducted analyses of variant calling results. RC, DAY, WM, CAC, TLR, JL all contributed to figure generation. RC, DAY, WM, MS, RM, JdI, ESM, CX, KDP all contributed writing to the manuscript. CAC and TLR helped facilitate the group meetings and coordinate the group’s efforts. RC led the group’s efforts. All author’s contributed to determining the direction of the research and evaluating analysis results.

---

Supplemental Figure 1. Indel Calls across the length of the SARS-CoV-2 Genome. In-frame calls are indicated in blue, frameshifting calls are in orange. Calls in homopolymer regions are indicated by “HM.” Many of the calls are made in homopolymer regions, those earlier in the genome are more likely to be frameshifting, and only a few positions have InDels called many times across the dataset considered.

Supplemental Figure 2. Agreement across pipelines with and without recommended parameters, recent dataset. A+B+C+D) Agreement across pipelines without recommended parameters. E+F+G+H) Agreement across pipelines with recommended parameters. A+E) Agreement on Illumina SNP calls. B+F) Agreement on Illumina InDel calls. C+G) Agreement on Oxford Nanopore (ONT) SNP calls. D+H) Agreement on ONT InDel Calls. For each figure, the bars indicate the number of variants called by the groups indicated by filled circles below, across the whole dataset.

Supplemental Figure 3. Difference in variant call frequencies across the length of the reference genome for each pipeline. A+B) Illumina platform data. C+D) ONT platform data. A+C) SNP calls. B+D) InDel Calls. For each pipeline, each row indicates a genomic position at which any pipeline called a variant. The color map indicates the difference in the frequency of the calls at that position, across the whole dataset, compared to the average frequency of calls made by all groups.

Supplemental Figure 4. Agreement across platforms with and without recommended parameters, recent dataset. A+B) Agreement between platforms without recommended parameters. C+D) Agreement between platforms with recommended parameters. A+C) Agreement between platforms on SNP calls. B+D) Agreement between platforms on InDel calls. For each figure, only those sample for which both Illumina and ONT platform data had at least one variant call that passed all the filters was considered. The total height is normalized to the total number of calls made by each pipeline, with the light blue portion indicating calls made on both platforms for a given sample, the medium blue indicating calls made only for the Illumina data, and the dark blue indicating calls made only for the ONT data.

