## Supplementary figures and images for "Towards increased accuracy and reproducibility in SARS-CoV-2 next generation sequence analysis for public health surveillance"

### Supplemental Figure 1

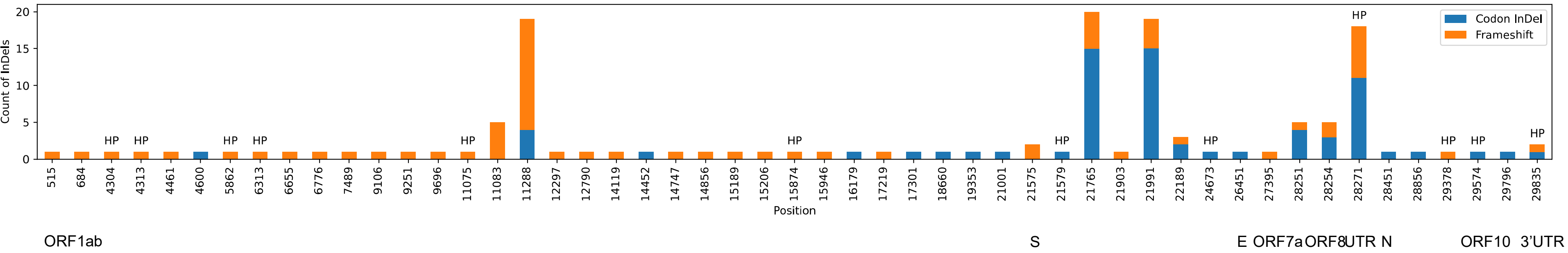

### Supplemental Figure 2

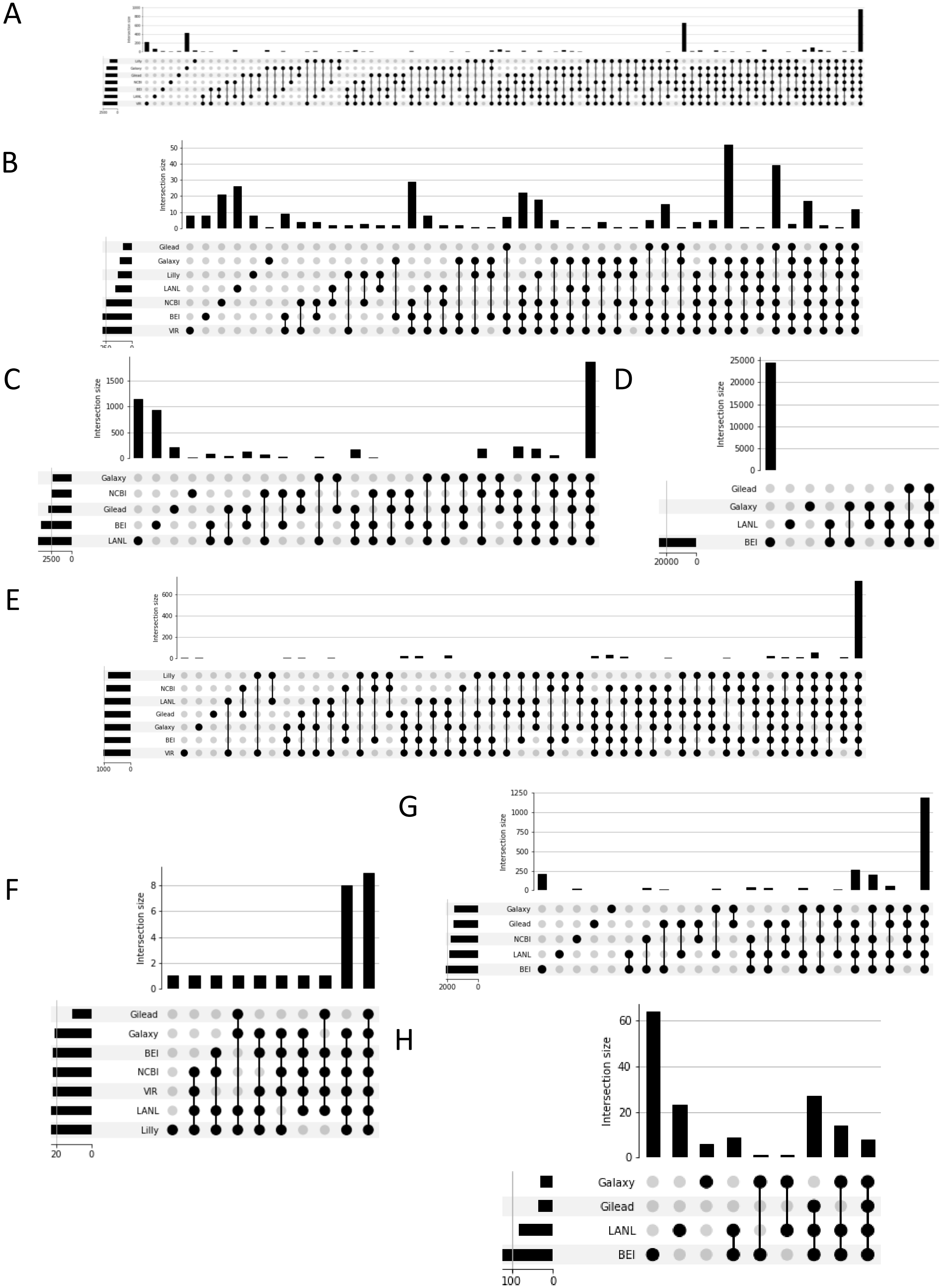

### Supplemental Figure 3

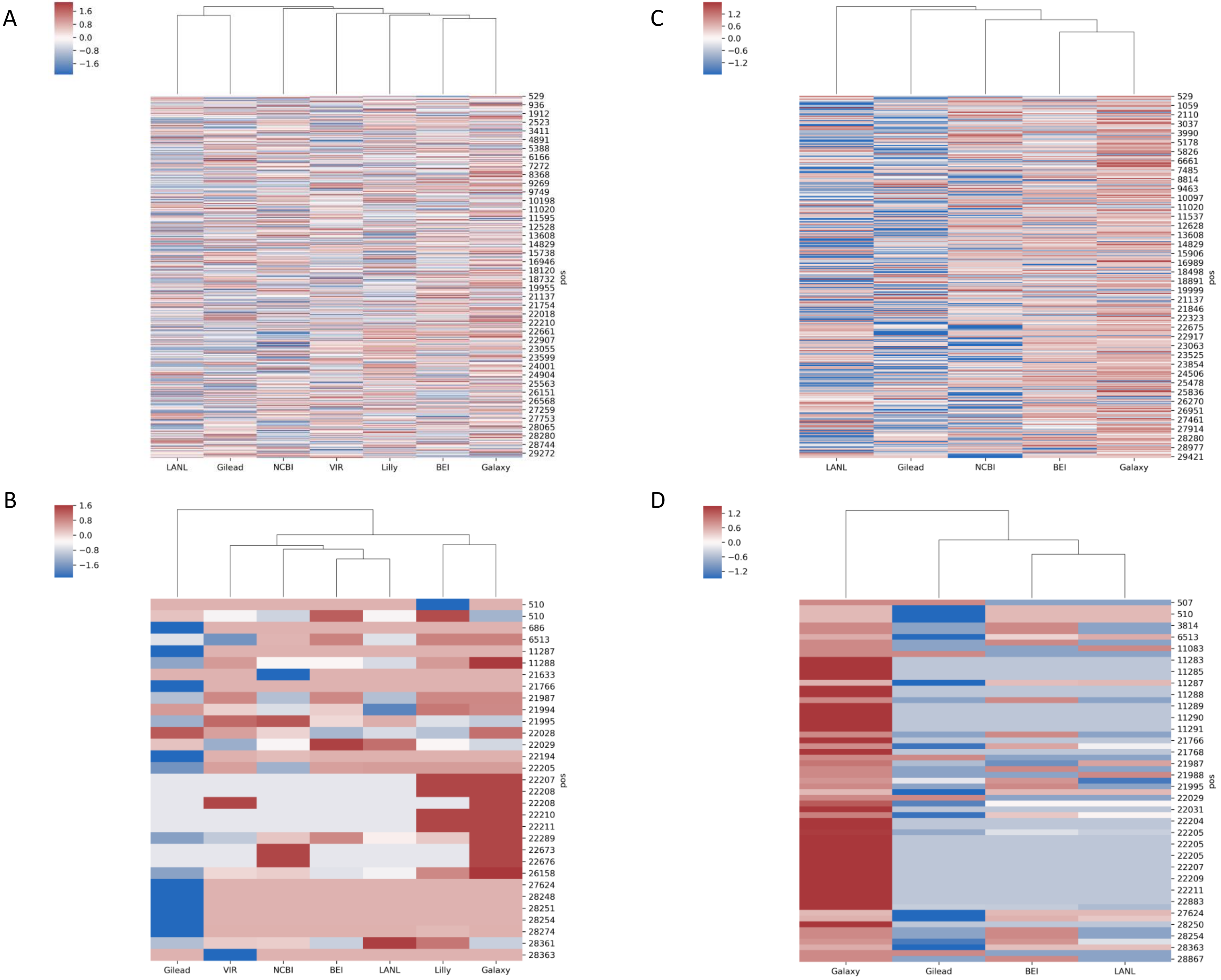

### Supplemental Figure 4

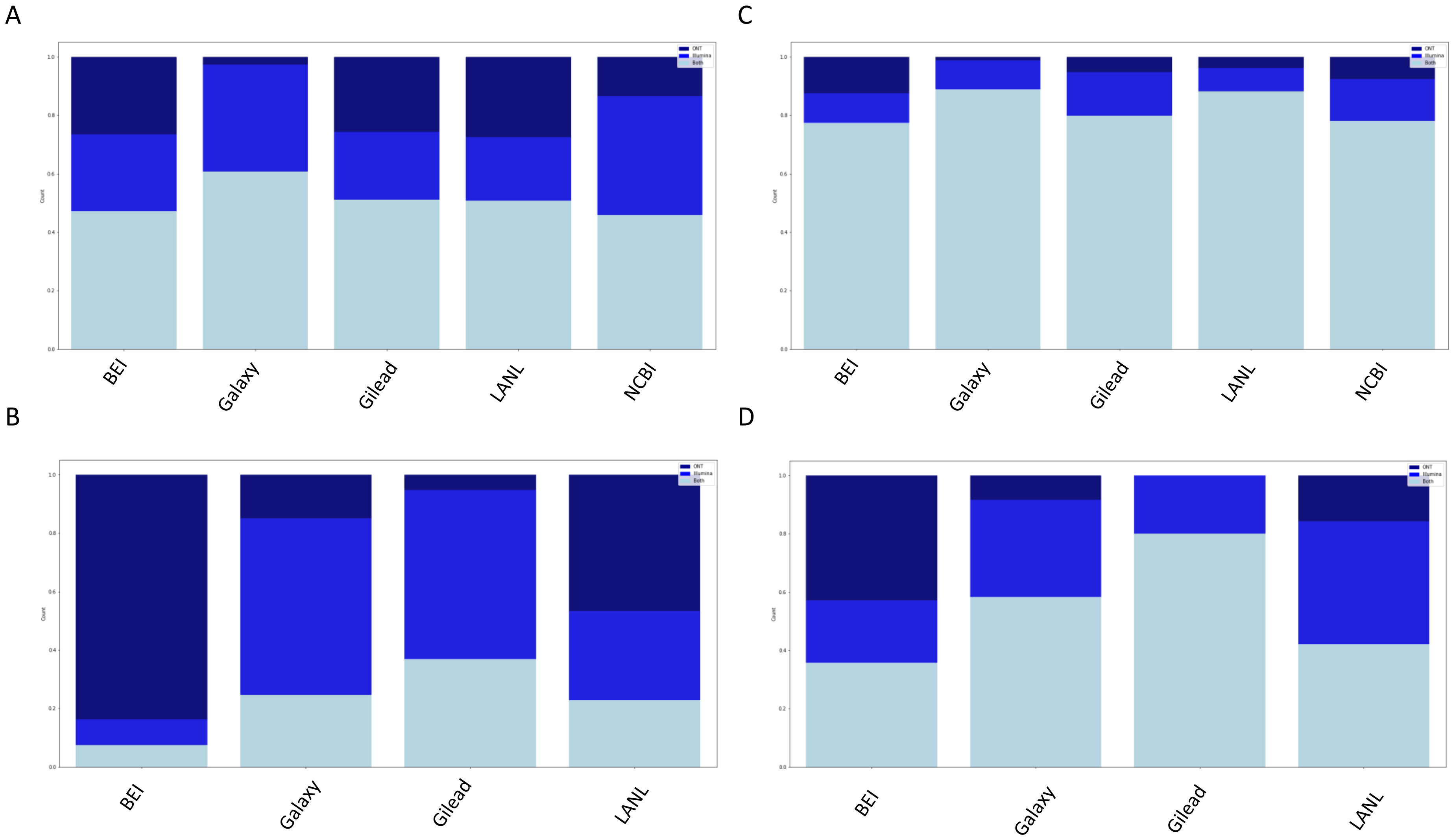
